# Autonomic Neuropathy Is Associated with An Increase in Type-1 Cytokines in People Living With HIV

**DOI:** 10.1101/2024.10.15.618447

**Authors:** Bridget R. Mueller, Mitali Mehta, Maya Campbell, Niyati Neupane, Gabriela Cedillo, Gina Lee, Kaitlyn Coyle, Jinging Qi, Zhihong Chen, Mary Catherine George, Jessica Robinson-Papp

**Affiliations:** Icahn School of Medicine at Mount Sinai, Department of Neurology; New York City, NY, USA; Icahn School of Medicine at Mount Sinai, Human Immune Monitoring Center (HIMC); New York City, NY, USA

**Keywords:** autonomic nervous system, dysautonomia, baroreflex, neuropathy, inflammation

## Abstract

**Purpose:** Pre-clinical studies have demonstrated direct influences of the autonomic nervous system (ANS) on the immune system. However, it remains unclear if ANS-immune connections delineated in pre-clinical studies underlie the relationship between autonomic dysregulation and chronic inflammatory diseases in patients with HIV. This study had three aims: 1.) Examine the relationship between IL-6 and the parasympathetic/vagal component of baroreflex sensitivity (BRS-V) in people with HIV; 2.) Determine if the subtype and severity of HIV-autonomic neuropathy (AN) would predict distinct immunotypes; 3.) Compare the burden of non-AIDS-related co-morbidities between immunotypes.

**Methods:** 79 adults with well-controlled HIV underwent a standard battery of autonomic function tests summarized as the Composite Autonomic Severity Score and vagal and adrenergic baroreflex sensitivity (BRS-V and BRS-A).^1^ Levels of immune biomarkers were measured in all participants using the Target 96 Inflammation Panel on the Olink proteomics platform and immunotypes were identified using unbiased, non-negative matrix factorization. Mass cytometry (CyTOF) was completed on a subset of participants with and without autonomic neuropathy (N = 10).

**Results:** Reduced BRS-V predicted higher levels of IL-6 (p=0.002). A pro-inflammatory immunotype defined by elevations in type 1 cytokines (IL-6, IL-17) and increased numbers of CD8+ T-cells was associated with autonomic neuropathy characterized by deficits in sympathetic nervous system activity (aOR=4.7, p=0.017). This pro-inflammatory immunotype was older with a greater burden of co-morbidities.

**Conclusion:** Deficits in the parasympathetic/cardiovagal and the sympathetic nervous system are associated with inflammation and disease burden in people living with HIV. Future longitudinal research is needed to examine causality.

## 1.1 Introduction

Preclinical studies have demonstrated the importance of the autonomic nervous system (ANS) in regulation of the innate and adaptive immune system. In humans, cardiovascular autonomic dysfunction has been demonstrated in patients with chronic inflammatory disorders including Long Covid (LC) syndrome ^2,3^, myalgic encephalomyelitis/chronic fatigue syndrome (ME/CFS)^4,5^, autoimmune disorders including multiple sclerosis,^6-8^, as well as rheumatoid arthritis.^9,10^ Decades of research have established that autonomic neuropathy (AN) is prevalent among people with HIV^11^ and HIV-AN is associated with significant morbidity and mortality ^12^ However, despite the significant prevalence of AN in HIV, the etiology of the comorbidity is unknown. Thus, a greater understanding of the ANS-immune network is needed in patients living with HIV.

In vitro and studies in animals have demonstrated that the extensive peripheral ANS, comprised of sympathetic efferents, parasympathetic (i.e. vagal) efferents, and sensory afferents has the ability to exert targeted, integrated, and rapid modulation of immune signaling through inflammatory reflexes.^13^ Lymphoid organs are highly innervated by sympathetic fibers and the sympathetic neurotransmitter norepinephrine (NE) regulates immune cell production and release of cytokines via α- and β-adrenergic receptors on located on immune cells. At high concentrations, NE released by sympathetic nervous system efferents has an inhibitory effect on immune cells through β-adrenergic receptors,^14^ while NE released by the sympathetic adrenal medullary (SAM) axis circulates at lower concentrations, preferentially activating pro-inflammatory, high-affinity α-receptors.^15,16^

The influence of parasympathetic/vagal activity is primarily anti-inflammatory and often referred to as the cholinergic anti-inflammatory pathway (CAP).^17^ Pre-clinical studies have established that activation of the CAP results in a reduction of inflammatory cytokine release, including interleukin-6 (IL-6), from splenic macrophages.^.18,19^ In vitro studies have demonstrated that the principal neurotransmitter of the parasympathetic nervous system, acetylcholine, released by vagal efferents, has anti-inflammatory effects on immune signaling. In a lipopolysaccharide-stimulated (LPS) human macrophage culture, the application of acetylcholine attenuated the release of pro-inflammatory TNFα, IL-6, and IL-18, but did not influence levels of the anti-inflammatory cytokine IL-10.^23^

Our knowledge of ANS regulation of the immune system in humans stems mainly from observational studies focused on parasympathetic/vagal function. Cardiac measurements indicative of lower resting vagal activity are present in patients with inflammatory bowel disease (IBD) as well as rheumatoid arthritis (RA) and neuropsychiatric conditions.^20,21^ In addition, vagal nerve stimulation has been associated with reduced levels of pro-inflammatory cytokines in patients with Sjogren’s syndrome and Crohn’s disease.^22,23^ Due to the complexity of the sympathetic nervous system and the methodological challenges associated with its measurement, the influence of sympathetic activity on immune signaling pathways in humans remain largely unexplored. This leaves a significant gap in our understanding of the ANS-immune network, as branches of the ANS do not act independently, but modulate immune signaling through an interconnected network.^13,18^

Our study had three goals. As pre-clinical studies have demonstrated the importance of IL-6 in the CAP^13^ and IL-6’s association with increased morbidity and mortality in people with HIV ^24,25^, we first investigated the relationship between IL-6 and the parasympathetic/vagal component of baroreflex sensitivity (BRS-V) in people with HIV. We chose to measure BRS-V because this reflex relies on caudal vagal circuitry common to the CAP. ^26^ Our second goal was to holistically examine the ANS-immune network and determine if the type (i.e. parasympathetic/vagal, cardiovascular sympathetic, and non-cardiovascular sympathetic) and severity of autonomic dysfunction was associated with distinct immune phenotypes identified by unsupervised non-negative matrix factorization (NMF) clustering analyses performed on a panel of 96 biomarkers of inflammation. Finally, as AN is associated with medical morbidity in people with HIV,^12^ we sought to compare the burden of non-AIDS-related co-morbidities between immunotypes. Given the availability of medications and devices that modulate both the ANS and the immune system, we hope a greater understanding of the ANS-immune network will rapidly translate into therapeutic studies.

## 2.0 Materials and Methods

### 2.1 Study Design and Patient Population

This is a cross-sectional observational study. Participants were recruited from a primary care clinic network within the Mount Sinai Health System in New York City. Eligible participants were identified by pre-screening clinic providers’ schedules and requesting approval from providers prior to contacting potential participants. Please see Supplemental Table 1 for full inclusion and exclusion criteria. Briefly, included participants were at least 18 years of age with a history of well-controlled HIV infection (plasma RNA load of ≤ 100 copies/ml for 3 months prior to enrollment) who were on a stable combination of antiretroviral treatment (CART) for at least 3 months. Patients with another diagnosis known to cause autonomic dysfunction (e.g., diabetes) not able to complete required autonomic testing (e.g., participant must be able to stand) or taking substances that impact the autonomic nervous system (e.g. urine drug testing positive for stimulants) were excluded. All procedures were performed in accordance with a protocol approved by the Institutional Review Board of the Icahn School of Medicine at Mount Sinai (ISMMS) and all participants provided written informed consent.

### 2.2 Autonomic testing procedures

Autonomic function tests (AFTs) are a standard battery of non-invasive tests (WRMedical®; https://wrmed.com/) which include sudomotor testing (QSWEAT), heart rate response to deep breathing, Valsalva maneuver (VM), and tilt table testing. The QSWEAT is performed by placing a capsule containing acetyl choline (ACh) on the skin in four standardized locations (forearm, proximal leg, distal leg, and foot). The capsule is attached to an automated system which delivers a small continuous electrical stimulus to the capsule causing iontophoresis of ACh into the skin, which triggers a reflexive sweat response collected by the capsule. The evoked sweat volume is measured and compared to standardized values. A non-invasive continuous beat-to-beat blood pressure (BP) monitoring device is attached to the participant’s finger and a 3-lead surface electrocardiogram and respiratory monitor are attached to the chest. BP, heart rate (HR), and respirations are recorded during the VM (forced exhalation to a pressure of 40 mmHg for 15 seconds), standardized paced deep breathing (HRDB), and a 10-minute head-up tilt test. It is important to note that while the QSWEAT is a direct measurement of sympathetic efferent function, cardiovascular responses to deep breathing and tilt table position changes are also influenced by other physiologic factors (e.g. arterial compliance, and activation of stress response systems). ^10^ Therefore, cardiovascular changes during autonomic testing are indirect measurements of parasympathetic or sympathetic activity.

### 2.3 Calculation of autonomic indices

The above-described procedures are used to calculate the Composite Autonomic Severity Score (CASS), which is a validated age- and sex-adjusted summary score reflecting overall autonomic function and is the sum of three sub-scores.^1^ The sudomotor (i.e. peripheral, non-cardiovascular sympathetic) sub-score uses data from the QSWEAT, the parasympathetic/vagal sub-score is based on changes in HR during deep breathing and VM, and the adrenergic (i.e., cardiovascular sympathetic) sub-score is based on BP changes during VM and tilt table testing. Given the medical complexity of our patient population, our lab uses the stringent threshold of a total CASS ≥3 to define AN; a sub-score of 1 defines mild dysfunction and a sub-score ≥2 identifies moderate to severe dysfunction.^27^

Baroreflex sensitivity (BRS) was calculated as previously described.^28^ Briefly, VM data is visually inspected by a trained, blinded technician. BRS-V is a measure of the compensatory cardiac response to a decrease in BP evoked during the forced expiration against a closed glottis and is calculated by dividing the change in RR interval during phase 2E of the VM by the change in systolic blood pressure. It is a continuous measurement expressed as milliseconds/mmHg. BRS-A is expressed in mmHg/second, and it is calculated by dividing the change in systolic blood pressure during phase 3 by the time required for SBP to recover following the release of VM.

### 2.4 Medical history and patient-reported outcome measures

Medical history and concomitant medications were obtained through participant interview and review of the electronic health record (EHR). To control potentially confounding influences of medications and comorbidities, several indices were included in multivariable analyses. First, as acetylcholine is a main neurotransmitter of the ANS, an anticholinergic burden (ACB) medication score was determined for all participants. Second, a Charlson Comorbidity Index was calculated (modified to exclude HIV/AIDS).^29-31^ Participants completed the Hospital Anxiety and Depression Scale (HADS)^32^ and the Perceived Stress Scale-14 (PSS-14).^33^

### 2.5 Biospecimen Collection and Proteomics Analysis

Whole blood was collected in EDTA tubes followed by isolation of peripheral blood mononuclear cell (PBMC) and plasma, as described previously.^34^ Serum was obtained concomitantly from serum-separator tubes. At our institution’s Human Immune Monitoring Center (HIMC), a shared core research facility, plasma IL-6 was measured as part of the Target 96 Inflammation Panel on the Olink proteomics platform (Uppsala, Sweden) as previously published.^34^

### 2.6 Cytometry by time of flight (CyTOF)

CyTOF was performed on PBMC samples from participants with AN (CASS ≥ 3) and without (CASS ≤ 1) selected purposively to maximize the difference in CASS between the groups while balancing sex and age (N = 5 per group) and without knowledge of cytokine profiles. Briefly, cell counts were performed on the Nexcelom Cellaca Automated Cell Counter (Nexcelom Biosciences) and cell viability was measured using Acridine Orange/Propidium Iodide viability staining reagent (Nexcelom). After washing cells in Cell Staining Buffer (CSB) (Fluidigm) Fc receptor blocking (Biolegend) and Rhodium-103 viability staining (Fluidigm) were performed simultaneously with surface markers for 30 minutes at room temperature. Cells were subsequently washed twice in CSB and then palladium barcoding was performed on each sample using the Cell-ID 20-Plex Pd Barcoding Kit (Fluidigm) following manufacturer’s instructions and pooled together. The pooled sample was fixed with 2.4% PFA (Electron Microscopy Sciences) followed by labeling with 125nM Iridium-193 (Fluidigm) and 2nM Osmium tetroxide (EMS) for 30 minutes at room temperature. Immediately prior to data acquisition, samples were resuspended at a concentration of 1 million cells per ml in Cell Acquisition Solution containing a 1:20 dilution of EQ Normalization beads (Fluidigm). The samples were acquired on a Helios Mass Cytometer (Fluidigm) equipped with a wide-bore sample injector at an event rate of <400 events per second. Prior to analysis, routine data normalization (Fluidigm software) and sample demultiplexing were undertaken.

### 2.7 Statistical analysis

All data were stored in REDCap. Descriptive statistics including frequencies, percents and medians with interquartile range were calculated as appropriate. Spearman rank correlation was performed to investigate the relationship between IL-6 and BRS-V. While CASS normative scoring is adjusted for age and sex, BRS is not. Therefore, multivariate logistic regression adjusting for age and sex was performed for analyses involving BRS. Kruskal-Wallis or Chi-square compared continuous and categorical variables between the four immunotypes, respectively. Bonferroni correction was applied to analyses involving multiple comparisons.

Immunotypes were defined using the R package for nonnegative matrix factorization (NMF) as previously described.^35^ Normalized protein expression of Olink inflammatory markers were compared between immunotypes. STRINGdb R package v2.16.4 was used for gene ontology (GO) enrichment analysis to elucidate key biological processes associated with immunotypes. The significance of GO terms was determined based on P-value (<0.001) and FDR (<0.05). To account for multiple testing and reduce the likelihood of false positives, we applied the Benjamini-Hochberg test for corrections. All analyses were conducted using SPSS version 28 and R 4.3.0.

## 3.0 Results

### 3.1 Study population demographics, clinical characteristics, and autonomic function

Participant demographics and medical characteristics are summarized in Table 1. Participants (N = 79) had an average age of 51.6 years (range: 25 – 73 years of age) and approximately three quarters were male. The majority had long-standing HIV with a self-reported mean of 22 years. African American was the most common race/ethnicity (48%), while 17% of participants were Hispanic/LatinX, 17% were non-Hispanic/LatinX white and 17% identified as other. With regard to medication, the mean ACB score for participants was 0.3 (SD = 0.8) with a range of 0 to 4. Regarding medical comorbidities, the median Charlson was 1.0 (IQR 0-1), which aligns with previously reported values in people living with HIV.^36^ Hypertension and hyperlipidemia were the most common comorbidities in our population (38.0% and 31.7% respectively; Table 1).

**Table 1:** Participant Demographics.

|  | Overall<br>N = 79 | Immunotype 1<br>N = 21 | Immunotype<br>2<br>N = 29 | Immunotype<br>3<br>N = 16 | Immunotype 4<br>N = 11 | p-value<sup>a</sup> |
| --- | --- | --- | --- | --- | --- | --- |
| Age, years* | 53.5 (42.0, 61.5) | 60.5<sup>b</sup><br>(52.8, 63.3) | 47.0 (18.5) | 52.5 (21.25) | 54.0 (27.0) | 0.043 |
| Sex, male | 58 (73.4) | 14 (66.7) | 22 (75.9) | 11 (68.8) | 8 (72.3) | 0.778 |
| Race ethnicity |  |  |  |  |  | 0.017 |
| African American | 38(48.1) | 6 (29.0) | 15 (51.7) | 14 (87.5)<sup>c</sup> | 2 (18.2) |  |
| Hispanic/LatinX | 14(17.7) | 7 (33.3) | 5 (17.2) | 0 (0) | 1 (9.1) |  |
| White | 13(16.5) | 4 (19.0) | 4 (13.8) | 1 (6.3) | 4 (36.4) |  |
| Other | 14(17.7) | 3 (14.3) | 5 (17.2) | 1 (6.3) | 3 (27.3) |  |
| Latest CD4+ count<br>(cells/mm<sup>3</sup>) | 578.0<br>(463.5,815.3) | 584.5<br>(244.0,837.5) | 640.5<br>(496.3, 851.3) | 547.0<br>(437.3, 855.3) | 521.0<br>(350.0, 758.0) | 0.496 |
| Time since HIV<br>diagnosis, years | 23.0<br>(15.0, 32.0) | 24.8<br>(19.4, 30.1) | 21.1<br>(17.3, 25.0) | 24.3<br>(19.2, 29.4) | 25.0<br>(19.4, 30.5) | 0.521 |
| HADS Anxiety | 7.1<br>(2.6, 11.6) | 7.6<br>(5.1,10.0) | 6.6<br>(5.1, 8.0) | 7.8<br>(5.2,10.5) | 7.0<br>(3.7, 10.3) | 0.818 |
| HADS Depression | 4.6<br>(0.81, 8.39) | 5.4<br>(3.5,7.2) | 4.1<br>(2.8,5.5) | 5.2<br>(3.2,7.1) | 3.8<br>(0.9, 6.5) | 0.541 |
| PSS-14 Score | 16.5<br>(9.0, 24.0) | 16.3<br>(12.5, 20.0) | 14.9<br>(12.4, 17.4) | 16.9<br>(12.3, 21.6) | 15.8<br>(9.4, 22.3) | 0.391 |
| CCI Score | 1.0 (0.0, 2.0) | 2.0 (1.0, 5.0)<sup>d</sup> | 0.5 (0.0, 2.0) | 1.0 (0.0, 2.0) | 1.0 (0.0, 6.0) | 0.013 |
| ACB Score | 0.3 (-0.5,1.1) | 0.7 (0.1 1.3) | 0.2 (0.0, 0.4) | 0.2 (0.0, 0.4) | 0.2 (0.0, 0.4) | 0.798 |
| Comorbidities |  |  |  |  |  |  |
| Hypertension | 30 (38.0) | 10 (47.6) | 13 (44.8) | 5 (31.3) | 2 (18.2) | 0.329 |
| Hyperlipidemia | 25 (31.65) | 9 (42.9) | 8 (27.6) | 5 (31.3) | 3 (27.3) | 0.570 |
| CAD | 3 (3.8) | 2 (9.5) | 1 (3.4) | 1 (6.3) | 0 (0) | 0.387 |
| Syphilis history | 19 (24.1) | 3 (14.3) | 11 (37.9) | 2 (12.5) | 3 (27.3) | 0.225 |
| HSV | 9 (11.4) | 4 (19.0) | 1 (3.4) | 1 (6.3) | 3 (27.3) | 0.086 |
| Asthma | 14 (17.7) | 4 (19.0) | 5 (17.3) | 2 (12.5) | 3 (27.3) | 0.819 |
| Obesity | 5 (6.3) | 3 (14.3) | 1 (3.4) | 1 (6.3) | 0 (0) | 0.308 |
| Osteoarthritis | 7 (8.9) | 4 (19.0) | 2 (6.8) | 0 (0.0) | 1 (9.1) | 0.209 |
| Solid tumor | 9 (11.4) | 3 (14.3) | 2 (6.8) | 1 (6.3) | 3 (27.3) | 0.262 |
| Unless otherwise stated, data are summarized as mean (95% confidence interval) or N (% of column).<br>*, median (interquartile range). |  |  |  |  |  |  |
| a, comparison across four immunotypes |  |  |  |  |  |  |
| b, p < 0.01 compared to immunotype 2 |  |  |  |  |  |  |
| c, p < 0.01 compared to immunotypes 1 and 4. |  |  |  |  |  |  |
| d, p < 0.05 compared to immunotype 2 |  |  |  |  |  |  |
| Abbreviations: CD4: Cluster of Differentiation 4; HADS: Hospital Anxiety and Depression Scale; PSS-14: Perceived Stress Scale; ACB: Anticholinergic Burden; CAD: Coronary Artery Disease; HSV: Herpes Simplex Virus |  |  |  |  |  |  |

76/79 participants completed autonomic nervous system testing. The total CASS ranged from 0-6 with a median score of 2.0. AN was common (44.7%) in our study population. Examination of CASS sub-scores showed that 42% of all participants had adrenergic (i.e., cardiovascular sympathetic) sub-score abnormalities, 57% had sudomotor (i.e. non-cardiovascular sympathetic) abnormalities, and 49% had cardiovagal (i.e., vagal/parasympathetic) dysfunction. In patients with AN, dysfunction across multiple domains of ANS function was common, with dysfunction in two domains present in 58.9% of participants and 29.4% demonstrating dysfunction in all three domains.

### 3.2 Increased IL-6 In Patients with Parasympathetic/Vagal Dysfunction

We assessed the relationship between IL-6 and parasympathetic/vagal function. We chose to measure IL-6 because elevated IL-6 has been shown to correlate with to increased morbidity and mortality in people with HIV 24,25 and pre-clinical studies have demonstrated IL-6 plays a key role in the CAP.^13^ Spearman rank correlation revealed a significant univariate correlation between IL-6 and BRS-V in the expected direction with reduced vagal function correlating to higher IL-6 (Spearman’s rho = -0.352, p=0.002; Figure 1). This was confirmed with multivariate, stepwise, linear regression with IL-6 as the outcome variable and BRS-V, age, ACB score, CD4+, sex, Charlson score, and diagnoses of anxiety and depression as the predictors; the final model retained BRS-V (p=0.012) and age (p=0.028).

**Figure 1:**
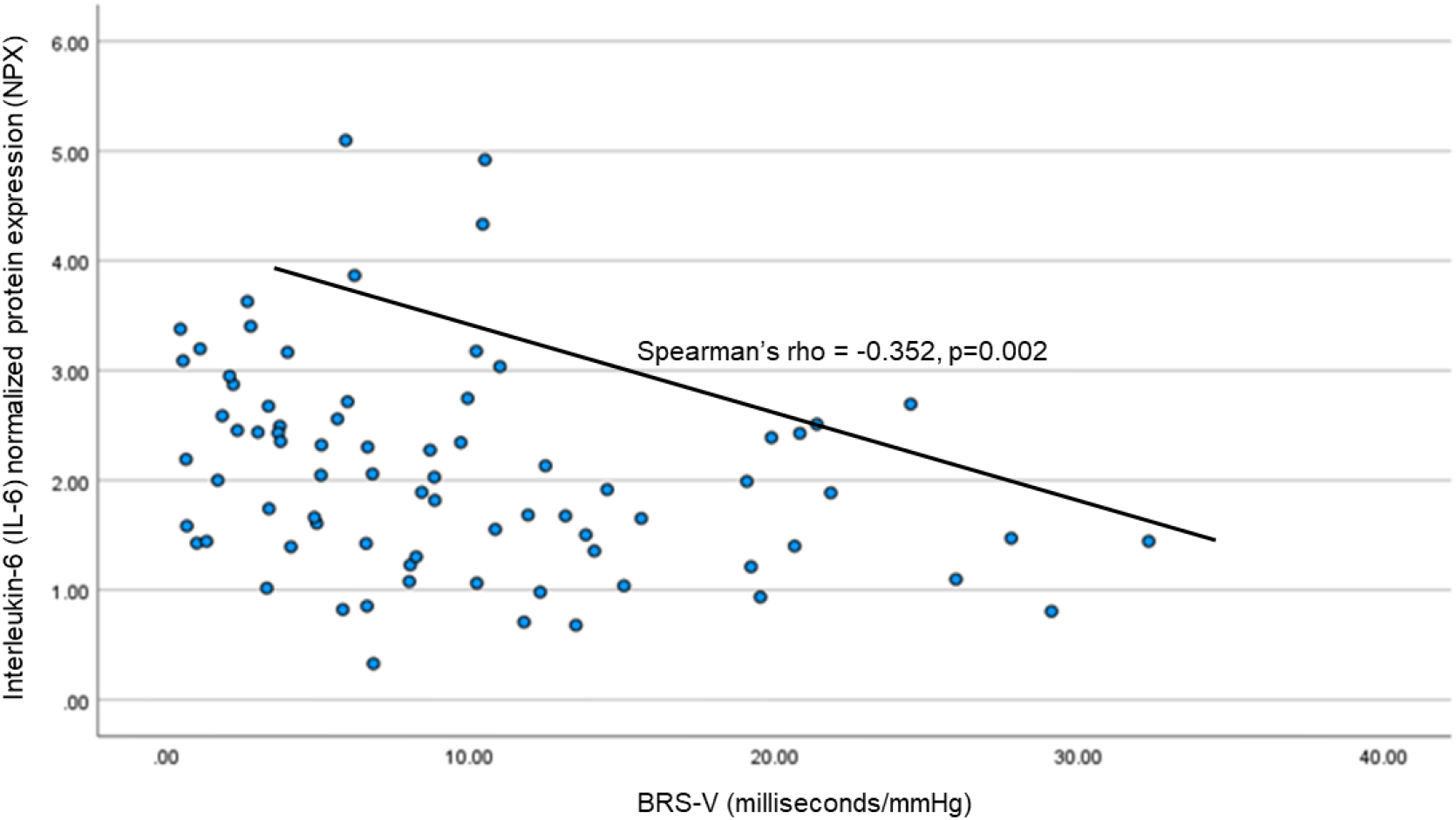
Lower vagal baroreflex sensitivity (BRS-V) is associated with higher plasma interleukin-(IL-6).

### 3.3 Unsupervised Clustering Identified a Pro-inflammatory Immunotype with Older Participants, and a High Prevalence of AN and Burden of Co-morbid Illness

Next, we assessed the profile of inflammatory proteins using unbiased, non-negative matrix factorization (NMF) and found four clusters or immunotypes, with distinct differentially expressed protein expression profiles, as demonstrated in the resulting heatmap (Figure 2A). These immunotypes differed with respect to prevalence and severity of AN, age, as well as burden of co-morbid illnesses (Tables 1 and 2). Of note, sex, CD4+ count, HADS scores, ACB score, and PSS-14 scores did not differ between the four immunotypes (Table 1).

**Table 2:** ANS activity of immunotypes.

|  | Overall<br>N = 76 | Immunotype<br>1<br>N = 20 | Immunotype<br>2<br>N = 29 | Immunotype<br>3<br>N = 16 | Immunotype<br>4<br>N = 11 | p-value |
| --- | --- | --- | --- | --- | --- | --- |
| Total CASS | 2.0<br>[1.0,3.0] | 3.5<br>[2.0, 4.0] | 2.0<br>[0.0, 3.0] | 1.0<br>[1.0, 2.0] | 2.0<br>[0.0, 3.0] | 0.005<sup>a</sup><br>< 0.001<sup>b</sup> |
| Autonomic neuropathy<br>(CASS ≥ 3) | 34 (44.7) | 14 (70.0) | 13 (44.8) | 3 (18.8) | 4 (36.4) | 0.020<sup>a</sup><br>< 0.001<sup>b</sup> |
| Sudomotor CASS | 1.0 [0.0,2.0] | 2.0 [0.0, 4.5] | 0.0 [0.0,2.0] | 1.0 [0.0, 2.0] | 0.0 [0.0, 2.0] | 0.265<sup>a</sup><br>0.048<sup>b</sup> |
| Adrenergic CASS | 0.0 [0.0,1.0] | 1.0<br>[0.3,1.8] | 0.0<br>[ 0.0, 0.0] | 0.0<br>[0.0, 1.0] | 0.0<br>[0.0, 1.0] | < 0.001<sup>a</sup><br>< 0.001<sup>b</sup> |
| Cardiovagal CASS | 0.0 [0.0,1.0] | 1.0<br>[0.0, 2.0] | 1.0<br>[0.0, 1.0] | 0.0<br>[0.0, 0.75] | 0.0<br>[0.0, 1.0] | 0.161<sup>a</sup><br>0.198<sup>b</sup> |
| BRS-V, ms/mmHg | 8.0<br>[0.0,17.7] | 3.9<br>[1.4, 17.5] | 8.4<br>[3.5, 13.3] | 8.1<br>[6.0, 14.3] | 8.0<br>[4.1, 13.8] | 0.504<sup>a</sup><br>0.190<sup>b</sup> |
| BRS-A, ms/mmHg | 13.1<br>[3.1,23.1] | 10.1<br>[0.0, 21.5] | 13.3<br>[8.0, 19.7] | 13.6<br>[8.7, 18.2] | 14.9<br>[11.3, 18.0] | 0.630<sup>a</sup><br>0.248<sup>b</sup> |
| Data presented is median [1q, 3q] or N (% of column)<br>Abbreviations: BRS-A: adrenergic baroreflex sensitivity; BRS-V: vagal baroreflex sensitivity<br>a, comparison across four immunotypes<br>b, binary comparison between immunotype 1 and immunotypes 2-4 |  |  |  |  |  |  |

**Figure 2.**
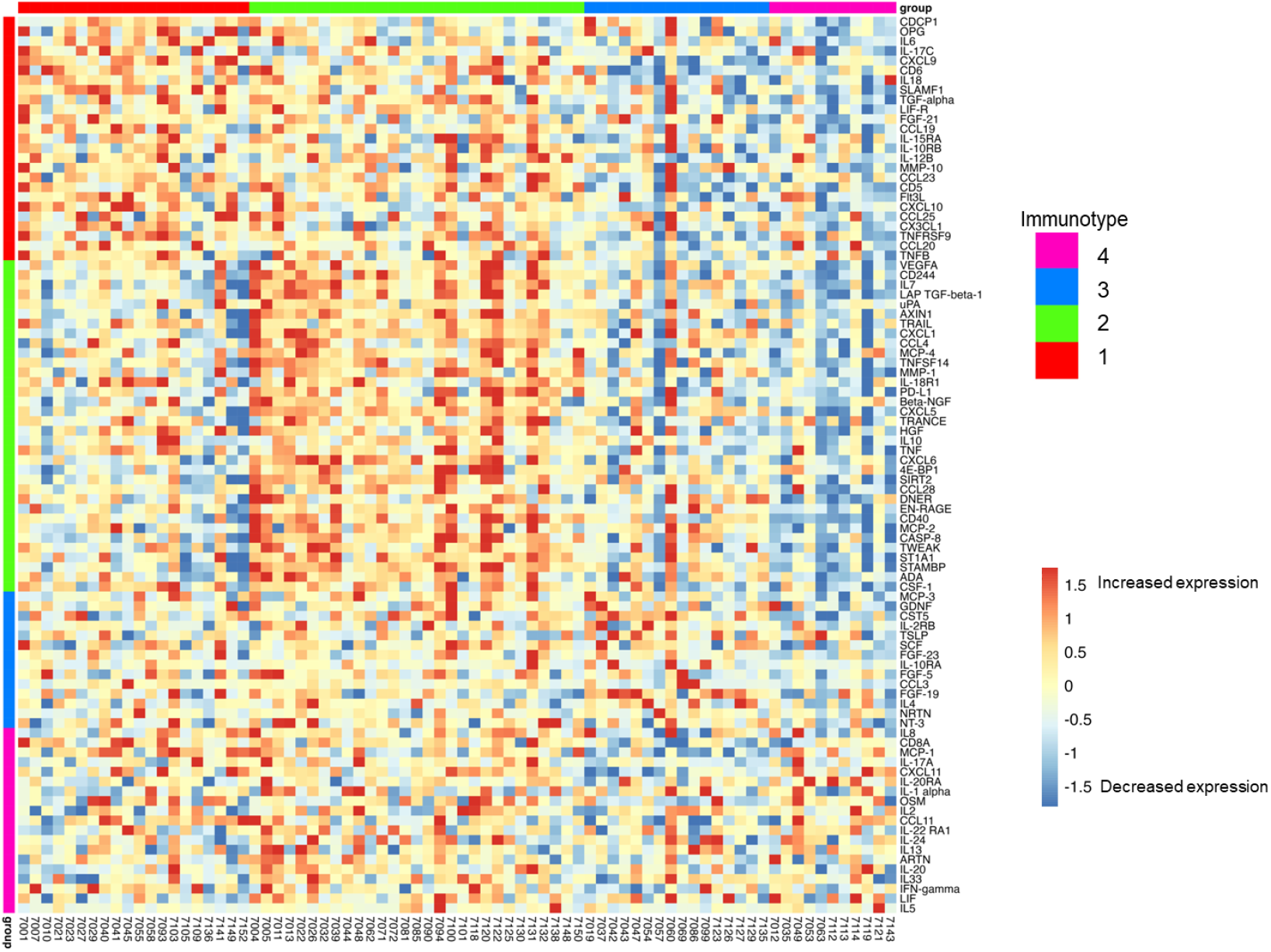
Heat map of 92 plasma immune biomarkers demonstrates four distinct immunotypes revealed by NMF clustering analysis and an elevation in type 1 cytokines in immunotype 1 (e.g. IL-6, IL-17) compared to other immunotypes (p < 0.05; see Supplemental Table 1) Immunotype 2 also exhibited increased pro-inflammatory markers, but at a lower level than immunotype 1, and showed an increase in anti-inflammatory markers compared to immunotype 1 (e.g. IL-10 p < 0.05). Colored vertical bars indicate the biomarkers whose elevation defined the four immunotype groups. Horizontal bars indicate the participants within each immunotype. Blue represents decreased relative protein expression while red represents increased expression.

<u>Immunotype 1</u> was defined by an upregulation of pro-inflammatory biomarkers including interleukins (IL-6, IL-17), and members of the TNF families (IL-18, TNF-β, TNFRSF9, OPG) which are regulated by the CAP. As shown in Figure 2, immunotype 1 was characterized by increased levels of interleukins involved in the expansion and activation of CD8+ T-cells, including IL-12B, IL-17c, and IL-10RB, and decreased levels of proteins that act to inhibit T-cell expansion including CASP-8.^37^ (See Supplemental Table 1 for normalized protein expression and pairwise comparisons across the four immunotype). GO analyses demonstrated an enrichment in proteins involved in T-helper 1 cell cytokine production, as well as the migration and chemotaxis of several immune cell subtypes (Figure 3A).

**Figure 3:**
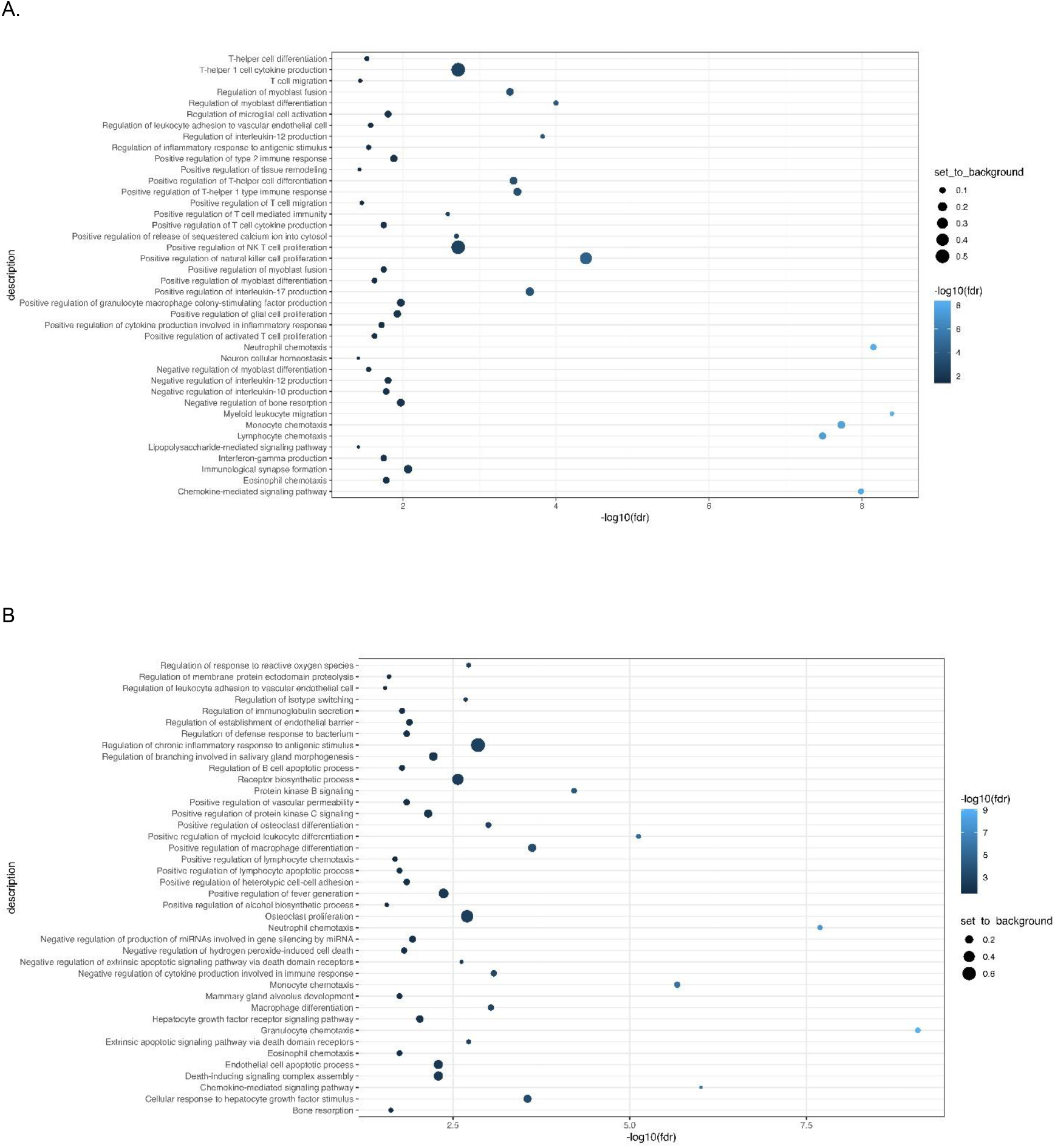
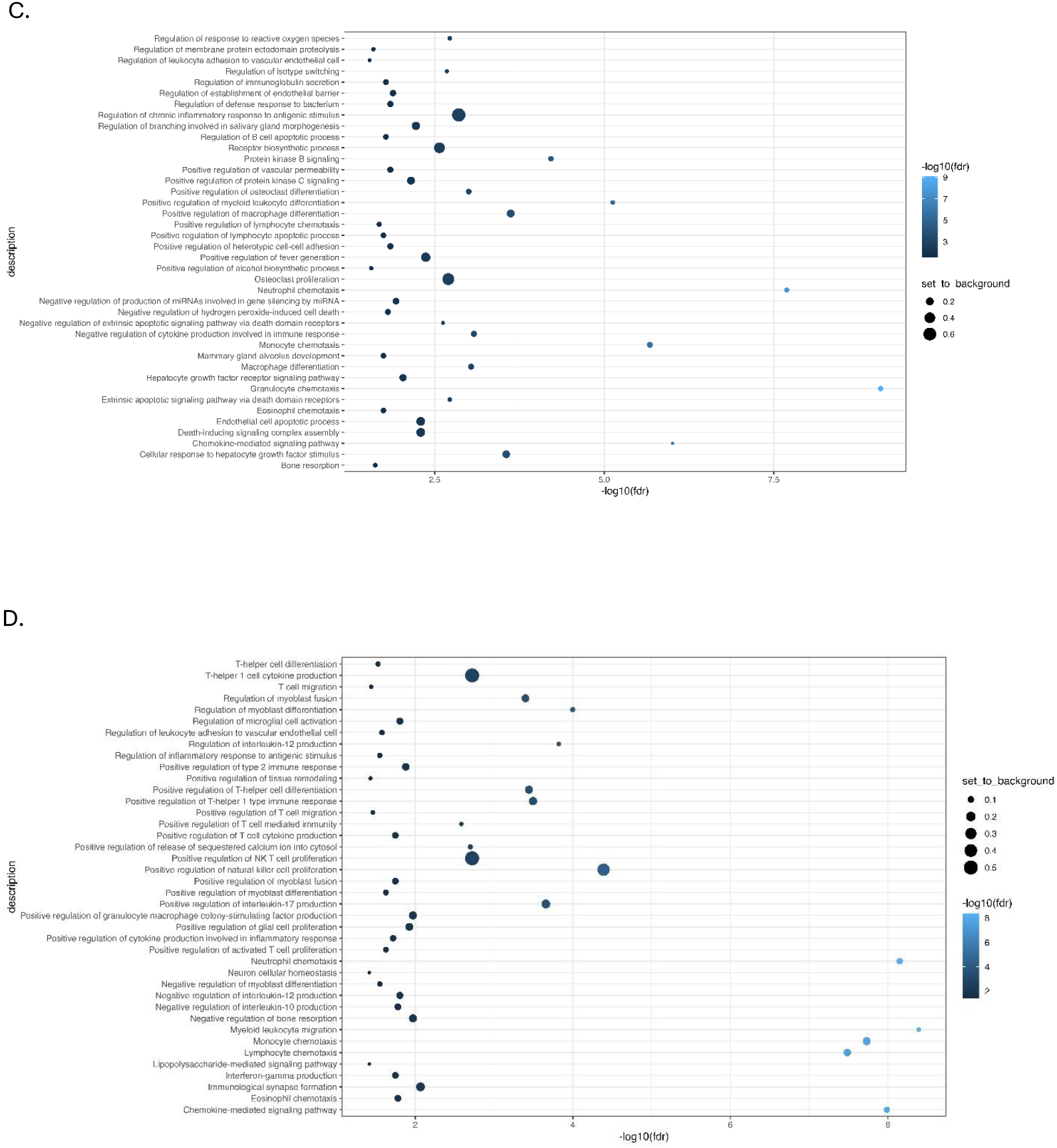
Gene Ontology (GO) analysis demonstrates increased expression of type 1 pro-inflammatory cytokines in immunotype 1 (A) and upregulation of proteins involved in the inhibition of pro-inflammatory cytokines, apoptosis, and immune signaling in immunotypes 2 (B) and 3 (C). Immunotype 4 (D) had an increased expression of proteins involved in intracellular signaling and immunoglobulin secretion.

Immunotype 1 was six years older than the study population overall (60.5 years versus 53.5 years) and had a higher median Charlson Comorbidity Index compared to other immunotypes. The most prevalent medical comorbidities were hypertension (47.6%) and hyperlipidemia (42.9%). Immunotype 1 also had the highest prevalence of AN (70%). In multivariate logistic regression participants with AN were almost five times more likely to be immunotype 1 than immunotypes 2-4 (aOR = 4.7, p=0.017) after adjusting for age, sex, ABC score, Charlson Index, and CD4+ count. All participants with moderate to severe AN (CASS > 4) were in immunotype 1 and no participants with a CASS of zero were in immunotype 1. Cardiovascular sympathetic nervous system deficits defined this immunotype as they were present in three-quarters of participants in immunotype 1, compared to just 20-30% in other immunotypes. (Table 2, Figure 4; p = 0.001)

**Figure 4.**
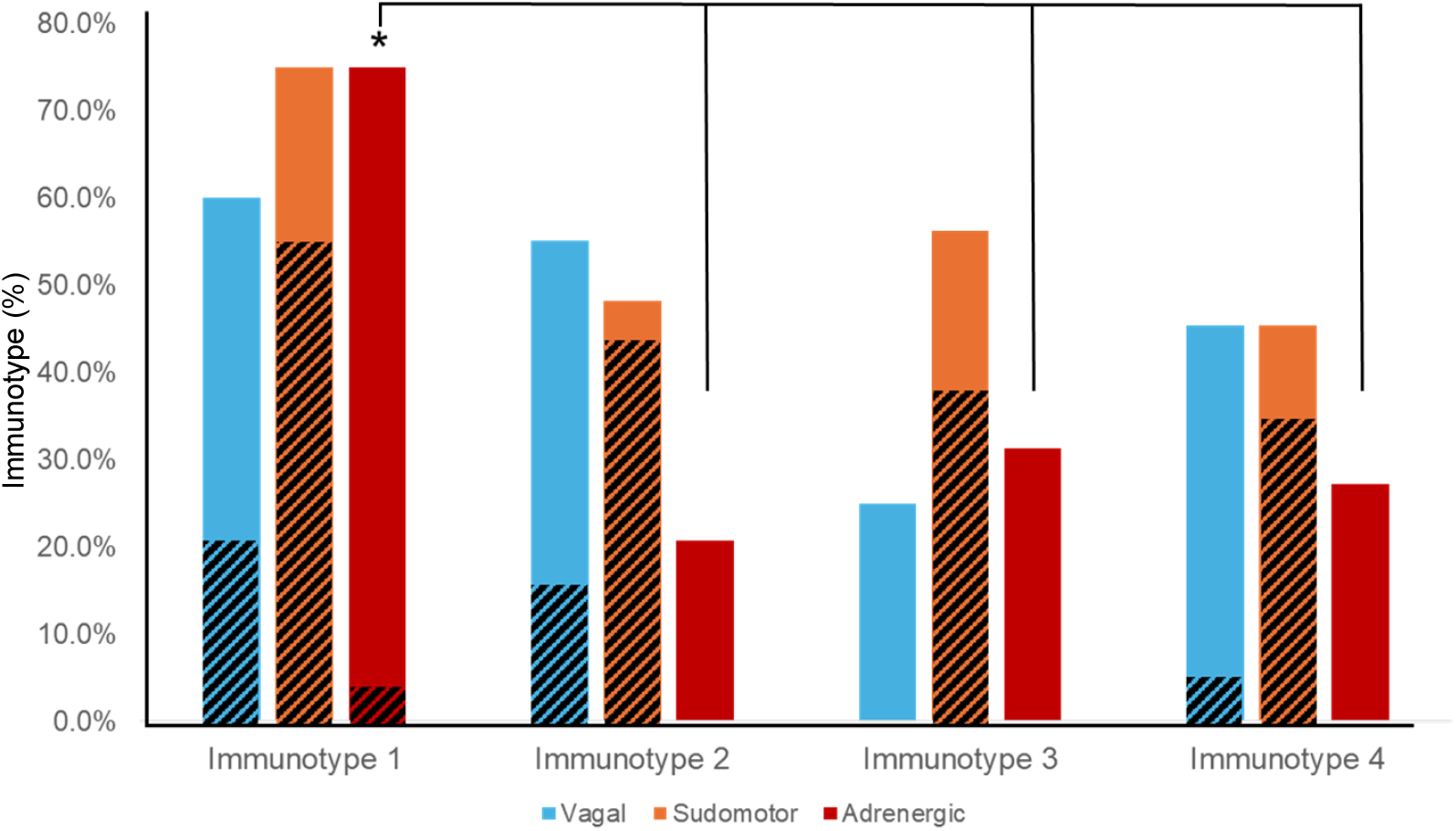
ANS testing revealed distinct patterns of parasympathetic/vagal, non-cardiovascular sympathetic (i.e. sudomotor), and cardiovascular sympathetic (i.e. adrenergic) deficits, as defined by a CASS sub-score ≥1, in immunotypes 1-4. Diagonal lines indicate prevalence of moderate-severe deficits (CASS sub-score ≥2) * p < 0.001 indicates a significantly higher prevalence of cardiovascular sympathetic (i.e. adrenergic) deficits in immunotype 1 compared to immunotypes 2, 3, and 4.

<u>Immunotype 2</u> also demonstrated a pro-inflammatory profile; however, the examination of normalized protein expression showed that many of the immunotype 1 defining proteins, including IL-18 and TNF-β, were upregulated in immunotype 2 to a lesser extent than in immunotype 1. The statistical significance of these differences was assessed using pairwise comparisons as listed in Supplemental Table 1. Similarly, while NMF analyses identified the pro-inflammatory vascular endothelial growth factor (VEGF), hepatocyte growth factor (HGF), IL-12, and Fms-related tyrosine kinase 3 ligand (FLT3LG) as immunotype 2 defining proteins, these proteins were also upregulated in immunotype 1, though to a lesser extent. Compared to immunotype 1, immunotype 2 demonstrated an increased level of *anti-* inflammatory cytokines and proteins indicative of cell apoptosis and the negative regulation of cytokine production (e.g. IL-10; p = 0.02 for pairwise comparison between immunotype 1 and 2 (Figure 2 and Supplemental Table 1)These results aligned with pathways identified in GO enrichment analyses (Figure 3B).

Immunotype 2 was significantly younger than immunotype 1 (47.0 versus 60.5 years) and had a significantly lower Charlson Comorbidity Index than immunotype 1 (0.5 versus 2.0). Immunotype 2 also had the shortest HIV disease duration, though this was not a statistically different result. Immunotype 2 had a lower prevalence of AN (44.8%) than immunotype 1, which was due to a reduced prevalence of adrenergic deficits (20% versus 75%).

<u>Immunotype 3</u> had a different immune signaling profile compared to immunotypes 1 and 2. The pro-inflammatory proteins elevated in immunotypes 1 and 2 were not elevated in immunotype 3 (Supplemental Table 1) and similar to immunotype 2, GO enrichment analysis demonstrated upregulation of proteins involved in the *negative* regulation of cytokine production and differentiation of immune cells (Figure 3C). Defining proteins identified by NMF included the glial derived protein family (GDNF and NTN), neurotrophin 3 (NTF3) and IL-4, a cytokine involved in the differentiation of TH cells into TH2 cells.^38^

Demographically, the age and comorbidity burden of immunotype 3 did not differ from the study population average. However, immunotype 3 had significantly more African Americans than other immunotypes (87.5%).

<u>Immunotype 4</u> also had a lower inflammatory profile compared to immunotypes 1 and 2, though there was significant upregulation of several pro-inflammatory proteins related to immune cell chemotaxis including CXCL11 and monocyte chemoattractant protein-1 (MCP-1), which distinguished it from immunotype 3. (Supplemental table 1). GO analysis identified enriched expression of pathways involved in intracellular signaling and positive regulation of immunoglobulin secretion (Figure 3D).

Immunotype 4 had the highest prevalence of solid tumors, HSV, and asthma compared to other immunotypes, although this was not statistically significant. Approximately one-third of participants in immunotype 4 had AN (36.4%) but, like immunotype 3, the majority of autonomic dysfunction was mild (Figure 4).

### 3.4 CyTOF

To gain a greater understanding of how AN alters the immune profile, we performed a cellular profile analysis using high-parameter CyTOF, comparing the cellular composition between patients diagnosed with and without AN. As described in the methods, participants were selected purposefully to maximize the difference in CASS, and without knowledge of their cytokine profiles. Interestingly, we found that all patients in the AN group belonged to immunotype 1. CyTOF results showed an increased abundance of CD8+ T-cells in patients with AN compared to those without AN, which was particularly pronounced for the terminally differentiated effector memory subtype of the CD8+ T cells (Figure 5).

**Figure 5.**
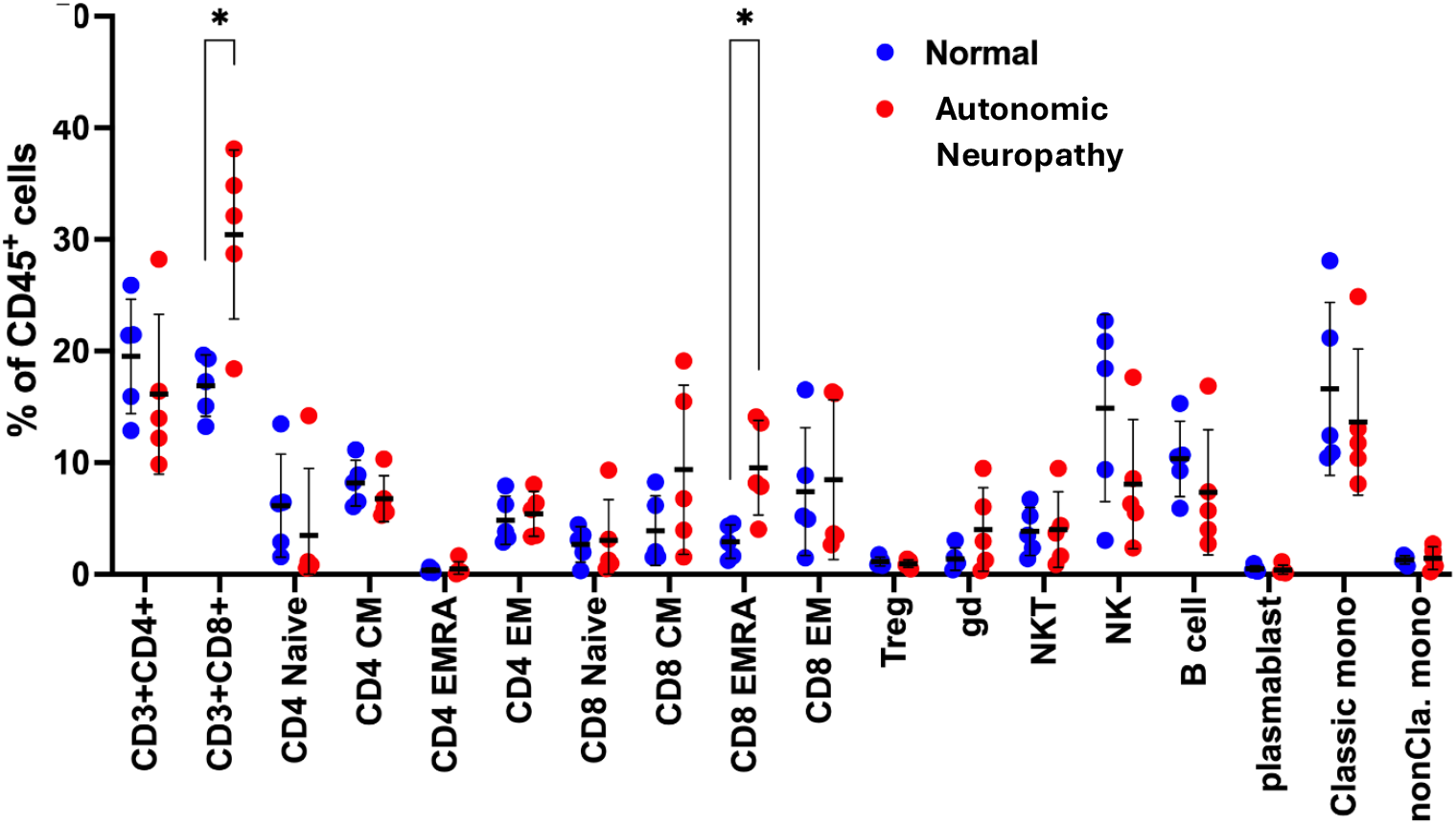
Comparison of immune cell type between participants with normal autonomic function and patients with autonomic neuropathy. **\*** p < 0.01 indicates significant increase in proportion of CD3+ CD8+ and CD8 EMRA (terminally differentiated effector memory T cell) sub-populations of T cells. Autonomic neuropathy is defined as CASS ≥ 3. Normal ANS function defined as CASS ≤ 1.

## 4.0 Discussion

We utilized comprehensive ANS testing and a broad-scale proteomic approach to assess the ANS-immune network in 79 people living with HIV. We found autonomic neuropathy (AN) was prevalent (44.7%) and predicted a pro-inflammatory phenotype characterized by elevations in type-1 cytokines, a predominance of CD8+ T-cells, and a higher burden of co-morbid illness in people living with HIV. These findings provide novel evidence for the clinical significance of parasympathetic/vagal and sympathetic pathways in the regulation of immune signaling.

This is the first study to demonstrate that diminished vagal baroreflex sensitivity, predicted elevated IL-6 in people with HIV. IL-6 plays a pivotal role in vascular inflammation, endothelial dysfunction, and adverse cardiovascular events.^39,40^ Yet, despite pre-clinical studies demonstrating IL-6’s regulation by the CAP, our understanding of this pathway in humans is limited. The majority of studies examining the relationship between parasympathetic/vagal activity and systemic inflammation use resting heart rate variability (HRV) measurements.^20,21,40,41^ However, resting HRV is associated with the activity of prefrontal-amygdala pathways ^42^and is decreased in stress-related mood disorders.^43^ Our own lab found reduced resting HRV in patients with normal peripheral vagal pathways.^44^ We found only one study that used a reflexive measure of vagal activity to assess the relationship between plasma IL-6 and caudal parasympathetic/vagal circuitry. In young patients with type I diabetes without AN, lower HRDB correlated to higher levels of plasma IL-6.^45^ Although cardiovagal baroreflex is influenced by arterial compliance^10^, and, thus, is an indirect measurement of vagal efferent activity, our results support the relevance of the CAP in patients with chronic inflammatory AN. Future studies will examine the relationship between IL-6 and measures of sympathetic/adrenergic function in patients living with HIV.

Unsupervised NMF clustering analysis revealed distinct immunotypes in people with HIV, supporting previous work that has demonstrated low and high inflammatory subgroups in people with HIV.^46^ Immunotype 1 was significantly older and had a higher burden of comorbid illness compared to immunotype 2. While both immunotypes 1 and 2 demonstrated upregulation of pro-inflammatory proteins associated with Type 1 T-cell expansion, immunotype 2 also had an enrichment of proteins involved in the *negative* regulation of inflammation and apoptosis, which was supported by GO analysis. The upregulation of these inhibitory proteins distinguished immunotype 2 from immunotype 1. The enrichment of CD8+ T-cells in patients with AN supports the proteomics analyses showing an upregulation of plasma immune biomarkers associated with proliferation of Type 1 T-cells in immunotype 1. The association of HIV-AN with a greater abundance of CD8+ T-cells is particularly relevant given that low CD4/CD8 ratio has been implicated in multiple poor outcomes in people with HIV including neurocognitive disorders, malignancy, and cardiovascular disease.^47^

Surprisingly, the pro-inflammatory immunotype 1 did not display a greater prevalence or severity of parasympathetic/vagal dysfunction. Instead, pathology in the sympathetic nervous system distinguished immunotype 1 from the younger and healthier immunotype 2. The increased prevalence of cardiovascular sympathetic deficits was particularly striking (75% in immunotype 1 versus 20% in immunotype 2). Non-cardiovascular sympathetic deficits, as measured by the sudomotor sub-score, were also more common in immunotype 1, indicating widespread sympathetic dysfunction. Pre-clinical studies provide an anatomic and physiologic basis of the anti-inflammatory impact of the sympathetic nervous system.^14^ In vitro studies have demonstrated that local release of NE from sympathetic efferents inhibits the production and release of pro-inflammatory cytokines through actions at β-adrenergic receptors located on the surface of immune cells.^14^ While we could not directly assess the sympathetic efferents innervating organs of the immune system, in more advanced HIV-AN with evidence of both cardiovascular and non-cardiovascular sympathetic impairment (as immunotype 1 displayed), it would be unlikely that they would be selectively spared.^48^ Moreover, loss of sympathetic innervation of lymph nodes has been demonstrated in Simian Immunodeficiency Virus (SIV) SIV models.^49^ Future studies should examine if organ-specific deficits in post-ganglionic sympathetic efferent activity are present in people living with HIV.

The significant age difference between immunotype 1 (median 60.5 years) and 2 (median 47.5 years) warrants discussion. Determining the influence of aging on HIV-AN progression may be confounded by the common correlation between age and duration of HIV disease and the fact that younger people with HIV did not experience the early HIV epidemic (i.e., lack of treatment followed by neurotoxic treatments). Interestingly, disease duration did not significantly differ between immunotypes 1 and 2 and suggests that aging may contribute to deficits in cardiovascular sympathetic activity that characterize immunotype 1. We hypothesize that with time, the autonomic, immune and clinical characteristics of immunotype 2 may evolve to resemble those of immunotype 1. Clinical and pre-clinical studies indicate aging comprises peripheral nerve structure and function through mechanisms that involve inflammation, and thus may be accelerated in the context of the pathologic inflammation of chronic HIV.^50,51^ Unmyelinated, type C fibers of the sympathetic nervous system may be more vulnerable to aging than type A and B vagal efferent projections.^52^ We hypothesize that dysfunction of sympathetic fibers leads to an increase in pro-inflammatory activity in patients living with HIV, however, longitudinal research examining the progression of HIV-AN and its influence on the immune network is needed to explore this hypothesis and establish causality.

The diminished prevalence of AN in immunotype 3 may have contributed to its lower inflammatory profile. However, it is important to note that a significantly greater proportion of immunotype 3 identified as African American (87.5%) and while isolating the biological influence of race on the immune system is complicated by cultural and psychosocial influences, there is evidence for race-based differences in immune signaling which may confound conclusions regarding the ANS’s influence on immune signaling.^53^ Similarly, immunotype 4 had a higher prevalence of solid tumors, HSV, and asthma, and thus, AN is unlikely to be the primary influence on immune signaling.

There are certain limitations to our study. First, the single center, cross-sectional design and a small sample size prohibit establishing mechanistic causation and larger, longitudinal studies are needed. However, this work does provide important support for a pre-clinical literature that demonstrates direct modulation of the immune system by the ANS. Second, while female representation in our study aligns with HIV disease prevalence, the lower enrollment makes it difficult to examine the likely influence of sex differences on ANS-immune function. Finally, as mentioned in the methods section, cardiovascular responses to deep breathing or changes in body position provide indirect information about parasympathetic and sympathetic responsivity, as they are also influenced by other physiological factors including vascular compliance. Although our study used the sudomotor test to measure sympathetic efferent activity, we did not employ additional direct tests of autonomic activity such as catecholamine plasma levels in response to standing or intraepidermal nerve fiber density on skin biopsy. Therefore, to acknowledge this limitation, we chose a higher CASS threshold to identify autonomic neuropathy.

In conclusion, our results provide important evidence that the cholinergic anti-inflammatory pathway (CAP), established by pre-clinical studies, exists in humans and is relevant to the pathology of chronic inflammatory disorders. In addition, these data demonstrate the importance of the sympathetic nervous system in the development of a pro-inflammatory immune signature associated with a higher burden of co-morbid disease and suggest that longitudinal examination of the time course of HIV-AN progression may provide important insight regarding the influence of aging on this process. Finally, our findings illustrate that a comprehensive evaluation of the ANS-immune network advances our understanding of core mechanisms that lead to immune system dysregulation associated with increased morbidity and mortality in people with HIV and provide a rationale for future research to develop therapeutic strategies focused on modulation of the sympathetic and parasympathetic/vagal nervous system.

## Acknowledgments

This work was supported by a grant from the National Institute of Diabetes and Digestive and Kidney Diseases (NIDDK; R01DK122853) and the National Institute of Health (NIH) Helping to End Addition Long Term (HEAL: K12NS130673). This work was also supported in part through the computational and data resources and staff expertise provided by Scientific Computing and Data at the Icahn School of Medicine at Mount Sinai and supported by the Clinical and Translational Science Awards (CTSA) grant UL1TR004419 from the National Center for Advancing Translational Sciences.

