## Supplemental Tables 1 & 2 for "Autonomic Neuropathy Is Associated with An Increase in Type-1 Cytokines in People Living With HIV"

| **Supplemental Table 1:** Normalized protein expression levels in immunotypes 1-4, with pairwise comparisons corrected for multiple comparisons. | | | | | | | | | | | | | |
| --- | --- | --- | --- | --- | --- | --- | --- | --- | --- | --- | --- | --- | --- |
| **Analyte** | **Group 1 median** | | **Group 2 median** | | **Group 3 median** | **Group 4 median** | ***Group 1 versus 2 p-value*** | ***Group 1 versus 3 p-value*** | | ***Group 1 versus 4 p-value*** | ***Group 2 versus 3 p-value*** | ***Group 2 versus 4 p-value*** | ***Group 3 versus 4 p-value*** |
| 4E-BP1 | 5.174 | | 5.881 | | 5.439 | 5.362 | 0.001 | | 0.633 | 0.169 | 0.017 | 0.100 | 0.824 |
| ADA | 5.289 | | 5.749 | | 5.626 | 4.967 | 0.035 | | 0.626 | 0.272 | 0.254 | 0.000 | 0.272 |
| ARTN | -0.266 | | -0.150 | | -0.376 | -0.281 | 0.489 | | 0.203 | 0.929 | 0.068 | 0.587 | 0.576 |
| AXIN1 | 5.407 | | 6.405 | | 5.869 | 5.151 | 0.000 | | 1.000 | 0.907 | 0.028 | 0.000 | 0.272 |
| Beta-NGF | 1.483 | | 1.610 | | 1.498 | 1.501 | 0.012 | | 0.230 | 0.828 | 0.050 | 0.029 | 0.875 |
| CASP-8 | 1.899 | | 2.926 | | 2.334 | 1.904 | 0.000 | | 0.023 | 0.814 | 0.017 | 0.000 | 0.287 |
| CCL11 | 7.518 | | 7.162 | | 6.948 | 7.363 | 0.164 | | 0.203 | 0.760 | 0.071 | 0.399 | 0.110 |
| CCL19 | 9.728 | | 9.701 | | 9.210 | 8.866 | 0.703 | | 0.015 | 0.019 | 0.095 | 0.001 | 0.272 |
| CCL20 | 7.251 | | 7.361 | | 6.842 | 6.626 | 0.908 | | 0.761 | 0.061 | 0.016 | 0.040 | 0.834 |
| CCL23 | 11.248 | | 11.195 | | 10.483 | 11.117 | 0.937 | | 0.626 | 0.246 | 0.001 | 0.261 | 0.110 |
| CCL25 | 5.643 | | 5.440 | | 5.525 | 4.867 | 0.787 | | 0.171 | 0.134 | 1.000 | 0.083 | 0.302 |
| CCL28 | 1.403 | | 1.433 | | 1.286 | 1.059 | 0.316 | | 1.000 | 0.033 | 0.136 | 0.004 | 0.188 |
| CCL3 | 5.139 | | 5.234 | | 4.759 | 4.656 | 0.937 | | 0.954 | 0.017 | 0.088 | 0.002 | 0.531 |
| CCL4 | 5.327 | | 6.009 | | 5.544 | 5.058 | 0.082 | | 0.672 | 0.336 | 0.056 | 0.005 | 0.272 |
| CD244 | 4.871 | | 5.293 | | 4.843 | 4.437 | 0.000 | | 0.015 | 0.033 | 0.008 | 0.000 | 0.248 |
| CD40 | 11.058 | | 11.996 | | 11.313 | 10.148 | 0.001 | | 0.002 | 0.023 | 0.010 | 0.000 | 0.015 |
| CD5 | 5.061 | | 5.142 | | 4.572 | 4.424 | 0.671 | | 0.046 | 0.033 | 0.002 | 0.001 | 0.875 |
| CD6 | 4.203 | | 4.250 | | 3.288 | 3.533 | 0.928 | | 0.388 | 0.058 | 0.000 | 0.011 | 0.248 |
| CD8A | 8.894 | | 9.014 | | 8.185 | 8.396 | 0.960 | | 0.271 | 0.099 | 0.016 | 0.027 | 0.875 |
| CDCP1 | 3.122 | | 2.605 | | 2.512 | 2.084 | 0.042 | | 0.271 | 0.004 | 0.814 | 0.080 | 0.165 |
| CSF-1 | 8.762 | | 8.708 | | 8.431 | 8.264 | 0.935 | | 0.203 | 0.044 | 0.017 | 0.003 | 0.481 |
| CST5 | 6.679 | | 6.937 | | 7.337 | 6.183 | 0.563 | | 0.008 | 0.044 | 0.299 | 0.001 | 0.005 |
| CX3CL1 | 3.497 | | 3.203 | | 3.174 | 2.971 | 0.339 | | 0.020 | 0.017 | 0.969 | 0.105 | 0.260 |
| CXCL1 | 8.666 | | 9.664 | | 9.147 | 8.135 | 0.000 | | 0.145 | 0.294 | 0.164 | 0.000 | 0.110 |
| CXCL10 | 8.328 | | 7.882 | | 7.250 | 7.367 | 0.069 | | 0.969 | 0.017 | 0.017 | 0.083 | 0.845 |
| CXCL11 | 8.617 | | 8.865 | | 7.828 | 9.190 | 0.794 | | 0.000 | 0.703 | 0.000 | 0.753 | 0.038 |
| CXCL5 | 11.072 | | 12.639 | | 12.068 | 9.645 | 0.000 | | 0.461 | 0.087 | 0.112 | 0.000 | 0.022 |
| CXCL6 | 7.157 | | 8.256 | | 7.109 | 7.066 | 0.000 | | 1.000 | 0.561 | 0.000 | 0.000 | 0.747 |
| **Analyte** | **Group 1 median** | | **Group 2 median** | | **Group 3 median** | **Group 4 median** | ***Group 1 versus 2 p-value*** | | ***Group 1 versus 3 p-value*** | ***Group 1 versus 4 p-value*** | ***Group 2 versus 3 p-value*** | ***Group 2 versus 4 p-value*** | ***Group 3 versus 4 p-value*** |
| CXCL9 | 7.264 | | 6.712 | | 5.894 | 6.552 | 0.056 | | 0.090 | 0.033 | 0.010 | 0.656 | 0.137 |
| DNER | 8.144 | | 8.309 | | 8.291 | 7.787 | 0.301 | | 0.090 | 0.118 | 0.963 | 0.006 | 0.110 |
| EN-RAGE | 2.904 | | 3.339 | | 2.677 | 2.073 | 0.089 | | 0.775 | 0.294 | 0.039 | 0.000 | 0.091 |
| FGF-19 | 7.348 | | 7.631 | | 8.501 | 7.425 | 0.870 | | 0.954 | 0.584 | 0.008 | 0.725 | 0.107 |
| FGF-21 | 4.770 | | 4.303 | | 3.831 | 3.377 | 0.427 | | 0.090 | 0.033 | 0.461 | 0.109 | 0.545 |
| FGF-23 | 0.788 | | 0.866 | | 0.741 | 0.381 | 0.960 | | 0.848 | 0.087 | 0.926 | 0.056 | 0.097 |
| FGF-5 | 0.766 | | 0.855 | | 0.694 | 0.748 | 0.526 | | 0.128 | 0.607 | 0.315 | 0.115 | 0.747 |
| Flt3L | 8.465 | | 8.362 | | 8.158 | 8.091 | 0.483 | | 0.087 | 0.158 | 0.268 | 0.399 | 0.865 |
| GDNF | 0.729 | | 0.814 | | 0.828 | 0.492 | 0.743 | | 0.282 | 0.306 | 0.994 | 0.109 | 0.356 |
| HGF | 7.509 | | 7.518 | | 7.037 | 6.871 | 0.937 | | 0.743 | 0.044 | 0.023 | 0.002 | 0.692 |
| IFN-gamma | 7.349 | | 6.890 | | 6.232 | 6.880 | 0.483 | | 0.848 | 0.231 | 0.050 | 0.587 | 0.434 |
| IL-1 alpha | -0.044 | | 0.041 | | 0.173 | 0.138 | 0.339 | | 0.667 | 0.790 | 0.855 | 0.602 | 0.692 |
| IL10 | 1.399 | | 1.997 | | 1.628 | 1.433 | 0.038 | | 0.020 | 0.465 | 0.091 | 0.007 | 0.481 |
| IL-10RA | -0.343 | | -0.327 | | -0.576 | -0.494 | 0.928 | | 0.602 | 0.436 | 0.645 | 0.216 | 0.952 |
| IL-10RB | 5.572 | | 5.604 | | 5.451 | 5.301 | 0.928 | | 0.102 | 0.087 | 0.605 | 0.038 | 0.545 |
| IL-12B | 4.762 | | 4.916 | | 3.929 | 4.321 | 0.563 | | 0.324 | 0.318 | 0.000 | 0.121 | 0.151 |
| IL13 | -0.741 | | -0.746 | | -0.709 | -0.666 | 0.728 | | 0.271 | 0.486 | 0.933 | 0.753 | 0.875 |
| IL-15RA | 0.275 | | 0.215 | | 0.060 | 0.303 | 0.884 | | 0.103 | 0.561 | 0.268 | 0.740 | 0.638 |
| IL-17A | -0.225 | | 0.095 | | -0.519 | -0.442 | 0.301 | | 0.090 | 0.907 | 0.010 | 0.255 | 0.545 |
| IL-17C | 1.498 | | 1.284 | | 1.037 | 0.675 | 0.178 | | 0.203 | 0.071 | 1.000 | 0.222 | 0.287 |
| IL18 | 9.135 | | 9.021 | | 8.279 | 8.492 | 0.935 | | 0.672 | 0.109 | 0.019 | 0.088 | 0.725 |
| IL-18R1 | 5.811 | | 5.810 | | 5.490 | 5.390 | 0.960 | | 0.923 | 0.109 | 0.009 | 0.023 | 0.875 |
| IL2 | -0.674 | | -0.567 | | -0.564 | -0.653 | 0.153 | | 0.672 | 0.387 | 0.969 | 0.915 | 0.834 |
| IL-20 | -0.395 | | -0.056 | | -0.269 | -0.237 | 0.006 | | 0.672 | 0.631 | 0.046 | 0.183 | 0.905 |
| IL-20RA | -0.283 | | -0.275 | | -0.401 | -0.012 | 0.519 | | 0.402 | 0.104 | 0.616 | 0.165 | 0.260 |
| IL-22 RA1 | 0.235 | | 0.778 | | 0.421 | 0.724 | 0.129 | | 0.090 | 0.387 | 0.436 | 0.747 | 0.607 |
| IL-24 | 0.638 | | 0.939 | | 0.805 | 1.025 | 0.264 | | 0.336 | 0.436 | 0.723 | 0.887 | 0.875 |
| IL-2RB | -0.250 | | -0.134 | | -0.218 | -0.308 | 0.316 | | 0.090 | 0.766 | 0.994 | 0.231 | 0.545 |
| IL33 | -0.016 | | 0.151 | | 0.192 | 0.119 | 0.035 | | 0.388 | 0.401 | 0.933 | 0.407 | 0.545 |
| IL4 | -0.640 | | -0.730 | | -0.441 | -0.974 | 0.934 | | 0.492 | 0.766 | 0.315 | 0.433 | 0.173 |
| IL5 | -1.003 | | -0.560 | | -0.980 | -0.869 | 0.043 | | 0.775 | 0.465 | 0.104 | 0.747 | 0.670 |
| **Analyte** | **Group 1 median** | | **Group 2 median** | | **Group 3 median** | **Group 4 median** | ***Group 1 versus 2 p-value*** | | ***Group 1 versus 3 p-value*** | ***Group 1 versus 4 p-value*** | ***Group 2 versus 3 p-value*** | ***Group 2 versus 4 p-value*** | ***Group 3 versus 4 p-value*** |
| IL6 | 2.409 | | 2.000 | | 1.966 | 1.443 | 0.316 | | 0.388 | 0.017 | 0.497 | 0.049 | 0.481 |
| IL7 | 0.479 | | 1.118 | | 0.771 | -0.017 | 0.001 | | 0.672 | 0.380 | 0.039 | 0.000 | 0.107 |
| IL8 | 5.002 | | 5.032 | | 4.619 | 4.693 | 0.663 | | 0.002 | 0.655 | 0.112 | 0.261 | 0.845 |
| LAP TGF-beta-1 | 5.420 | | 5.779 | | 5.397 | 4.835 | 0.001 | | 0.206 | 0.061 | 0.010 | 0.000 | 0.061 |
| LIF | -1.031 | | -0.830 | | -0.928 | -0.792 | 0.025 | | 0.819 | 0.294 | 0.436 | 0.915 | 0.865 |
| LIF-R | 2.364 | | 2.424 | | 2.248 | 2.077 | 0.743 | | 0.388 | 0.017 | 0.436 | 0.002 | 0.097 |
| MCP-1 | 11.367 | | 11.053 | | 10.390 | 11.095 | 0.249 | | 0.775 | 0.401 | 0.009 | 0.671 | 0.022 |
| MCP-2 | 8.178 | | 8.938 | | 8.588 | 7.586 | 0.072 | | 0.492 | 0.044 | 0.497 | 0.000 | 0.005 |
| MCP-3 | 0.752 | | 0.918 | | 0.969 | 0.561 | 0.181 | | 0.508 | 0.306 | 0.524 | 0.006 | 0.151 |
| MCP-4 | 14.838 | | 15.176 | | 14.601 | 14.532 | 0.071 | | 0.775 | 0.436 | 0.003 | 0.007 | 0.875 |
| MMP-1 | 13.292 | | 14.241 | | 13.549 | 12.546 | 0.042 | | 0.633 | 0.095 | 0.058 | 0.000 | 0.110 |
| MMP-10 | 7.809 | | 7.659 | | 8.062 | 7.176 | 0.819 | | 0.206 | 0.044 | 0.196 | 0.043 | 0.028 |
| NRTN | -0.913 | | -0.807 | | -0.779 | -0.813 | 0.439 | | 0.619 | 0.679 | 0.933 | 0.952 | 0.981 |
| NT-3 | 2.042 | | 2.171 | | 2.201 | 1.832 | 0.291 | | 0.848 | 0.318 | 0.713 | 0.049 | 0.302 |
| OPG | 9.245 | | 9.041 | | 9.015 | 8.598 | 0.512 | | 0.171 | 0.058 | 0.994 | 0.029 | 0.272 |
| OSM | 2.846 | | 2.814 | | 2.095 | 2.811 | 0.960 | | 0.090 | 0.828 | 0.039 | 0.686 | 0.107 |
| PD-L1 | 4.967 | | 5.615 | | 4.943 | 4.816 | 0.012 | | 0.045 | 0.087 | 0.010 | 0.000 | 0.531 |
| SCF | 8.048 | | 8.144 | | 8.067 | 7.573 | 0.877 | | 0.090 | 0.294 | 0.969 | 0.029 | 0.151 |
| SIRT2 | 2.974 | | 4.340 | | 3.556 | 2.288 | 0.000 | | 0.013 | 0.465 | 0.040 | 0.000 | 0.048 |
| SLAMF1 | 1.221 | | 0.999 | | 0.894 | 0.568 | 0.012 | | 0.848 | 0.002 | 0.845 | 0.012 | 0.173 |
| ST1A1 | 3.993 | | 4.748 | | 3.966 | 3.399 | 0.010 | | 0.312 | 0.104 | 0.013 | 0.000 | 0.097 |
| STAMBP | 5.246 | | 6.302 | | 5.852 | 4.514 | 0.000 | | 0.383 | 0.457 | 0.010 | 0.000 | 0.038 |
| TGF-alpha | 2.161 | | 2.164 | | 1.925 | 1.781 | 0.826 | | 0.271 | 0.018 | 0.008 | 0.001 | 0.434 |
| TNF | 2.672 | | 2.570 | | 2.180 | 2.275 | 0.935 | | 0.848 | 0.099 | 0.010 | 0.029 | 0.952 |
| TNFB | 3.522 | | 3.363 | | 3.174 | 3.275 | 0.433 | | 0.383 | 0.231 | 0.323 | 0.617 | 0.834 |
| TNFRSF9 | 5.229 | | 4.998 | | 4.440 | 4.983 | 0.696 | | 0.388 | 0.272 | 0.007 | 0.407 | 0.272 |
| TNFSF14 | 3.789 | | 4.430 | | 3.725 | 3.179 | 0.003 | | 0.089 | 0.124 | 0.010 | 0.000 | 0.381 |
| TRAIL | 7.670 | | 7.712 | | 7.257 | 7.559 | 0.656 | | 0.854 | 0.387 | 0.046 | 0.080 | 0.845 |
| TRANCE | 3.685 | | 4.030 | | 3.368 | 3.776 | 0.197 | | 0.108 | 1.000 | 0.010 | 0.222 | 0.545 |
| TSLP | -0.007 | | -0.184 | | 0.251 | 0.157 | 0.563 | | 0.854 | 0.766 | 0.046 | 0.420 | 0.545 |
| TWEAK | 8.225 | | 8.470 | | 8.086 | 7.994 | 0.068 | | 0.667 | 0.066 | 0.017 | 0.000 | 0.151 |
| **Analyte** | **Group 1 median** | **Group 2 median** | | **Group 3 median** | | **Group 4 median** | ***Group 1 versus 2 p-value*** | | ***Group 1 versus 3 p-value*** | ***Group 1 versus 4 p-value*** | ***Group 2 versus 3 p-value*** | ***Group 2 versus 4 p-value*** | ***Group 3 versus 4 p-value*** |
| uPA | 9.811 | | 9.943 | | 9.551 | 9.583 | 0.175 | | 0.103 | 0.261 | 0.039 | 0.017 | 0.952 |
| VEGFA | 10.820 | | 10.976 | | 10.616 | 10.583 | 0.193 | | 0.122 | 0.124 | 0.019 | 0.006 | 0.511 |

**Supplemental Table 2**: Inclusion Criteria

Inclusion Criteria

- Age ≥18 years old
- Documentation of HIV-1 infection
- Stable combined anti-retroviral treatment (CART) for ≥3 months
- HIV viral load ≤100 copies/ml (within 3m)
- Willing to refrain from nicotine use for 24h prior to all testing;
- No contraindication to autonomic testing (e.g. uncontrolled glaucoma, heart rate not under sinus control)
- Ability to stand unassisted
- Urine test negative for stimulants and opiates/opioids
- No past or current medical condition known to cause autonomic dysfunction other than HIV (e.g. diabetes, Parkinson’s disease)
- No treatment history of chemotherapy or radiation known to cause autonomic dysfunction.
- Not taking medications with significant autonomic effects (e.g. sympathomimetics)
- No antibiotics within two weeks of testing
- Negative pregnancy test (if applicable)
